# Denuded peptidoglycan oligosaccharides enable the biochemical investigation of bacterial cell wall recognition, modification, and degradation

**DOI:** 10.64898/2026.07.08.737370

**Authors:** Benjamin G. Emmanuel, Grace DelMistro, Alexander C. Anderson, Chris Vandenende, Anthony J. Clarke, David Sychantha

## Abstract

Peptidoglycan is an essential component of the bacterial cell wall, providing mechanical strength and maintaining cell shape. It consists of glycan chains crosslinked by short peptide stems, resulting in a chemically heterogeneous macromolecule that remains challenging to study in a well-defined form. Access to discrete peptidoglycan fragments has therefore been critical for advancing biochemical and structural studies of cell wall-active enzymes. However, current synthetic, semi-synthetic, and cell wall extraction approaches remain limited by the complexity of carbohydrate chemistry and the difficulty of isolating pure, well-defined material. Here, we report a facile enzymatic approach for generating defined, denuded peptidoglycan oligosaccharides from the cell walls of two *Staphylococcus* specie*s*. These oligosaccharides, which terminate in *N*-acetylglucosamine and range from two to five disaccharide units in length, serve as substrates for a diverse panel of peptidoglycan-active enzymes that cleave or chemically modify the glycan backbone. We further show that these denuded oligosaccharides can be used in lysozyme-catalyzed transglycosylation reactions to generate *p*-nitrophenyl derivatives, enabling continuous colorimetric monitoring of peptidoglycan-cleaving enzymes. This method provides a practical route to defined peptidoglycan glycans and establishes a platform for further structural diversification, including stem peptide reattachment, quantitative enzyme assays, and structural characterization of peptidoglycan-binding proteins.

## Introduction

Bacterial cells are surrounded by a cell wall composed of a peptidoglycan (PG) layer that provides structural support for growth, division, and morphogenesis. These processes rely on PG to counteract the internal osmotic pressure exerted on the cell membrane, allowing bacteria to survive in hypotonic environments **(Fig. 1A)**. The essential role of PG in maintaining bacterial cell integrity has made it an enduring antibiotic target, as inhibiting PG biosynthetic enzymes is selectively toxic to bacteria. β-lactam and glycopeptide antibiotics are inhibitors of PG biosynthesis and have been in clinical use for over 50 years. However, due to the emergence of antibiotic resistance, interest in PG metabolism as a source of targets for the discovery of new antibiotics has been revived in recent years ^1^. This interest in PG extends further to its diverse roles as a major microbe-associated molecular pattern and a modulator of innate immunity^2^. PG mediates acute and chronic inflammatory responses impacting host physiology, behaviour, and drug interactions. Together, these roles have created a need to define how PG structure governs its biological functions, yet this remains difficult because PG is chemically heterogeneous, insoluble, and poorly suited for many conventional biochemical assays^3^.

**Figure 1.**
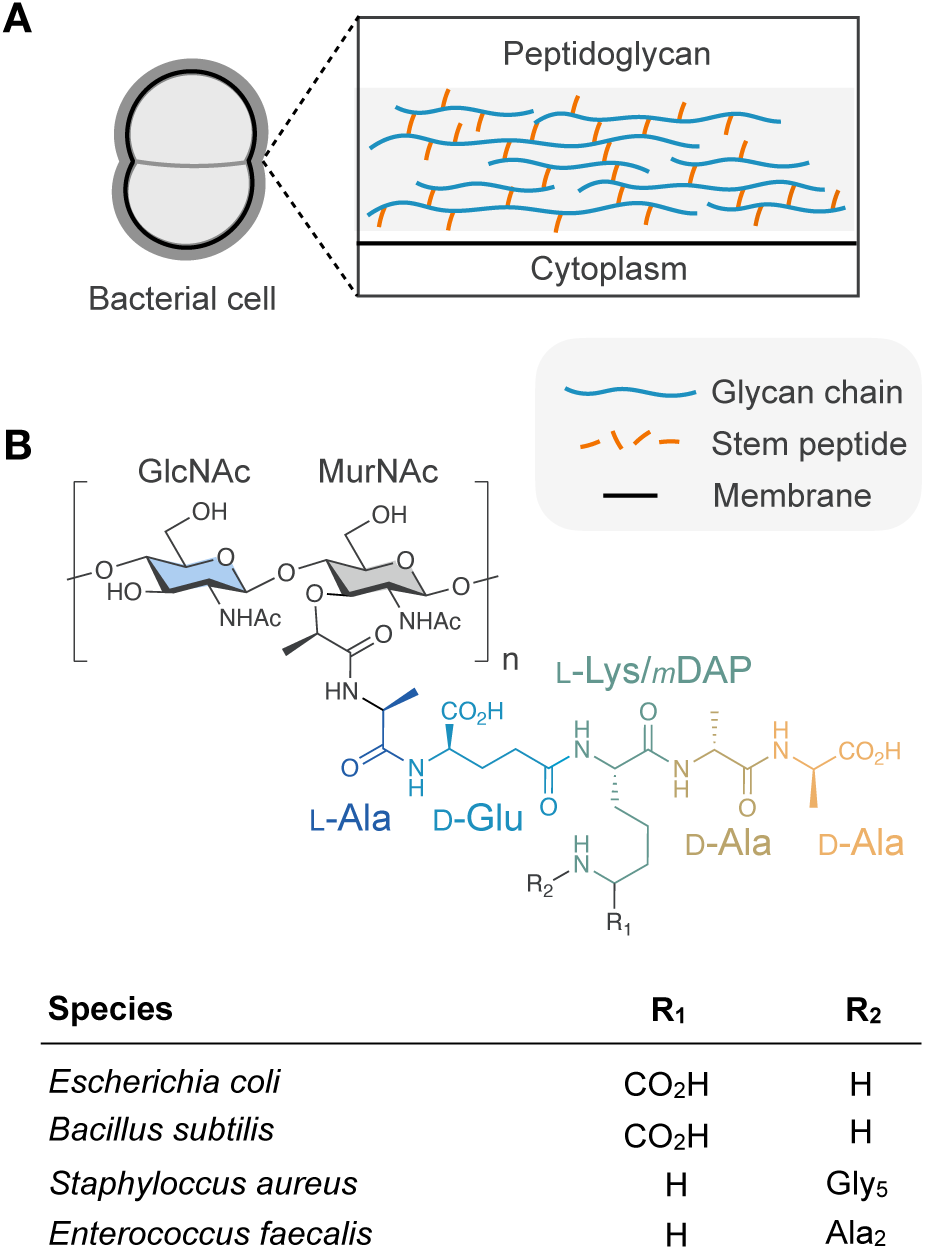
Overview of peptidoglycan structure. *A*, Simplified cartoon depiction of the arrangement of the cell wall around the bacterial cell membrane. *B*, Chemical structure of peptidoglycan, highlighting the stem peptide variability. Species-specific stem peptide structures denoted by *R* are defined in the corresponding table.

A major challenge to studying PG stems from its inherent physicochemical properties and heterogeneity. PG structure consists of conserved glycan chains of repeating β(1,4)-linked *N*-acetylglucosamine (NAG) and *N*-acetylmuramic acid (NAM) disaccharide units **(Fig. 1B)**^4^. The length of the glycan chain can range from 2 to greater than 30 units. Within the chain, NAM is further decorated with peptide stems that form crosslinks between adjacent chains, whose composition is species-specific **(Fig. 1B)**. The degree of crosslinking also varies among bacterial species and may approach 80-90% of the total peptide content in species with high degrees of crosslinking, such as staphylococci^5^. Altogether, the crosslinked glycans create PG’s three-dimensional structure, a single megadalton-sized insoluble molecule known as the sacculus **(Fig. 1A)**. This renders it resistant to standard biochemical and structural methods, which often require homogeneous, soluble ligands that are not commercially available. Accordingly, there is a strong interest in chemically defined, soluble PG fragments that can support the mechanistic studies of PG homeostasis and host interactions^3^.

Since the early 2000s, chemical synthesis methods for generating defined PG oligosaccharides have been instrumental in advancing understanding of PG biochemistry^6–16^. However, these approaches remain challenging, laborious, costly, and reliant on specialized synthetic expertise, which has limited their accessibility and broader adoption across the field^17^. To accelerate advances in PG biochemistry, we developed a practical, preparative-scale method to produce PG oligosaccharides directly from bacterial cell walls. Inspired by classical enzymatic digestion and fragment isolation methods developed by Sharon and Jeanloz in 1965^18^, our method enables rapid and practical milligram-scale production of well-defined PG oligosaccharides for functional assays and structural biology. The PG chains produced using our approach are derived from staphylococci, denuded (i.e., lacking a peptide stem), and terminate with a reducing NAG residue. Moreover, we demonstrate that these denuded glycans are suitable substrates for a diverse panel of PG-active enzymes, enabling further tailoring of these glycans. Finally, we leveraged these denuded glycans to produce a colorimetric substrate by attaching *p*-nitrophenol (pNP) to the reducing end of a denuded PG tetrasaccharide. When the cleavage of this derivative is coupled to an exo-β-*N*-acetylglucosaminidase, pNP is released, allowing continuous measurement of tetrasaccharide cleavage.

## Results

### Generating defined-length denuded PG chains through limited endo-β-*N*-acetylglucosaminidase digestion of staphylococcal *PG*

Historically, *Micrococcus luteus* has been used as a common source of PG starting material^18–22^. While this non-pathogenic monoderm soil bacterium produces a thick PG layer and has simple growth requirements, it exhibits slow growth rates and inconsistent culture yields in our laboratory.

Consequently, we explored alternative sources of PG and identified *Staphylococcus carnosus* and *Staphylococcus aureus* as suitable hosts for consistent large-scale cell wall preparation. These species were chosen for their simple growth requirements, fast growth rates, and the natural abundance of relatively short PG glycan chains^23^. Moreover, an advantage of using *S. carnosus* is that it is a non-pathogenic organism that may be safely grown on a large multi-litre scale.

The cell walls from these species were extracted with hot SDS and then treated with 0.5 M NaOH to remove covalently bound wall teichoic acids and any O-acetyl groups. Using this PG as the starting material, denuded PG chains with a degree of polymerization (DP) of 2, 3, 4, and 5 were obtained by controlled enzymatic digestion **(Fig. 2A)**. To achieve this, the PG stem peptide was first completely hydrolyzed from NAM by the recombinant amidase domain of *S. aureus* AtlA. The resulting free peptide products were protonated by adjusting the pH to 3.0, then adsorbed onto AmberChrom 5WX4 cation-exchange resin. The anionic PG glycans remained unbound and were conveniently separated from the resin by centrifugation. Typical yields were ∼500 mg of crude glycans from 20 L of culture.

**Figure 2.**
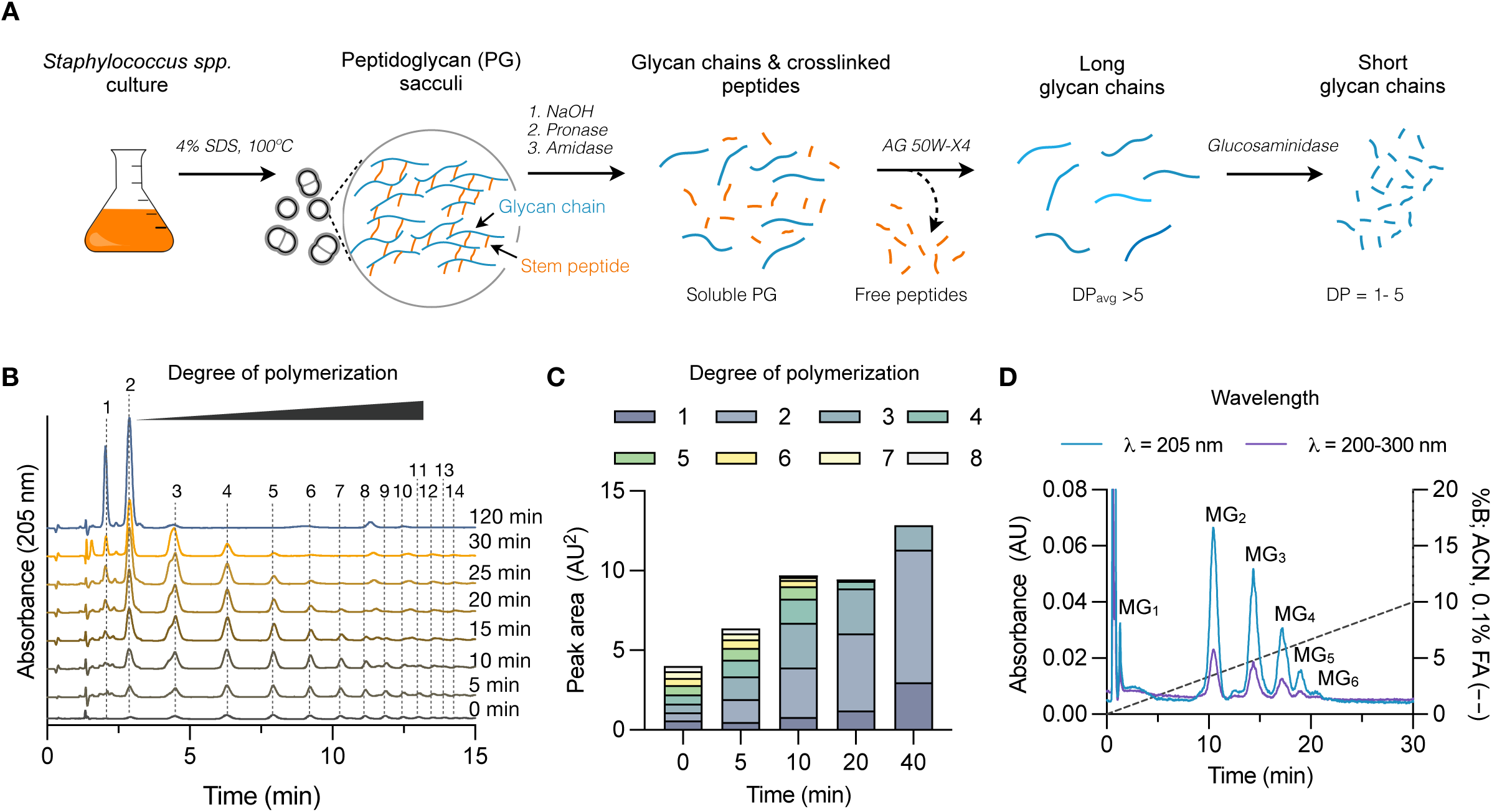
Limited digestion of *staphylococcal cell walls produces defined denuded PG chains.* *A,* Schematic workflow for the production of denuded glycan chains. *B*, HILIC analysis of a glycan chain digestion time-course by LytD compared to the initial glycan chain profile (0 min, black). The degree of polymerization is shown with dashed lines and corresponds to the elution time. *C,* Change in peak area of selected glycan chains over time. Peak area was obtained from a time-course of LytD-catalyzed glycan chain digestion, analyzed by HILIC. *D,* Flash chromatography chromatogram of large-scale glycan chain isolation following limited LytD digestion for 30 min.

Previous analysis of *S. aureus* glycan chains showed that they terminate with a reducing NAG residue, due to prominent β-*N*-acetylglucosaminidase activity^23^. Leveraging the existing abundance of reducing NAG-termini within the PG, the glycan mixture was subjected to time-limited (or enzyme-concentration-varied) digestion with the recombinant endo-β-*N*-acetylglucosaminidase LytD from *Bacillus subtilis*, as it produces a reducing NAG terminus post-cleavage. This produced well-defined products with little heterogeneity in oligosaccharide sequence **(Fig. 2B and Fig. S1 and S2)**. These denuded PG chains were denoted NAM-NAG_n_ (MG_n_), where *n* indicates the DP. From a starting glycan chain concentration of 30-100 mg/mL, hydrophilic interaction liquid chromatography (HILIC) analysis showed that LytD (20 μg/mL) converted glycan chains to oligosaccharides MG_2–5_ within 30 minutes **(Fig. 2C)**. If desired, significantly greater accumulation of MG_1_ and MG_2_ was achieved by extending the LytD digestion to 120 min or increasing the enzyme concentration, coincident with the conversion of oligosaccharides with a DP >3. HILIC analysis also detected a distribution of minor secondary species resulting from LytD digestion of *S. aureus* PG that were absent from *S. carnosus* PG. These products were considerably more polar than the denuded glycans and eluted later in the chromatogram, between 10 and 15 min **(Figure 2B)**. MS/MS analyses of these species gave masses consistent with denuded oligosaccharides (DP 2-5), with the C6-OH of NAM covalently attached to a single wall teichoic acid linker unit, ManNAc-(β1,4)-GlcNAc, via a phosphodiester bond **(Fig. S3)**. These fragments likely arose from incomplete NaOH hydrolysis of the insoluble PG starting material.

To purify each oligosaccharide, the mixture was separated by reversed-phase flash chromatography on a polar-end-capped C18 column, yielding well-resolved peaks of varying degrees of polymerization **(Fig. 2D)**. MS/MS confirmed the sequence of each glycan, indicating that oligosaccharides eluted in order of DP, with MG_2_ eluting first, followed by MG_3_, MG_4_, and MG_5_ **(Fig. S4 and S1).** After pooling and drying each pure fraction, each DP was ≥90% pure, as judged by HPLC.

### Denuded PG chains are suitable substrates for diverse glycan backbone-cleaving enzymes

To evaluate denuded glycans as enzyme substrates, we tested the MG_3_ hexasaccharide against thirteen PG backbone-cleaving enzymes with distinct cleavage-site specificities, reaction chemistries, and processivities. Overall, eleven lytic transglycosylases (LTs) from *Pseudomonas aeruginosa* as well as exo-muramidase and exo-glucosaminidase from *B. subtilis* were evaluated. Muramidases and LTs cleave the NAM–NAG linkage, producing reducing-end NAM and 1,6-anhydro-NAM (aM), respectively.

Whereas glucosaminidases target the NAG–NAM linkage, generating reducing-end NAG. MG_3_ was chosen for these assays as its length and sequence restrict the number of potential cleavage products, simplifying analysis. Given the order of the sugar subunits in MG_3_ and the specificity of LTs, it was anticipated that they would cleave directly in the middle, producing MGaM and GMG **(Fig. 3B)**. LC-MS analysis showed that five of the eleven LTs converted MG_3_ into the expected trisaccharide products, GMG and MGaM **(Fig. 3A)**. Among these LTs, SltB1, MltA, and RlpA were the most active with the MG_3_ substrate, converting most of it to MGaM. Notably, GaM and MGMGaM were also detected as additional products in SltB1 and MltA-catalyzed reactions, suggesting these enzymes remove the terminal NAM from MGaM as well as NAM or NAG of either end of MG_3_, consistent with their previously reported exo-activity **(Fig. 3B)**^9,24,25^. The other active LTs, Slt70 and MltD, converted less than 25% of MG_3_ into the trisaccharide products, indicating that they are significantly less active towards denuded glycans. The remaining enzymes, MltF, MltF2, MltG, SltB2, SltB3, and MltB, were not found to cleave MG_3_, suggesting that the absence of the stem peptide, glycan sequence, or structure of the non-reducing terminal sugar affects substrate binding.

**Figure 3.**
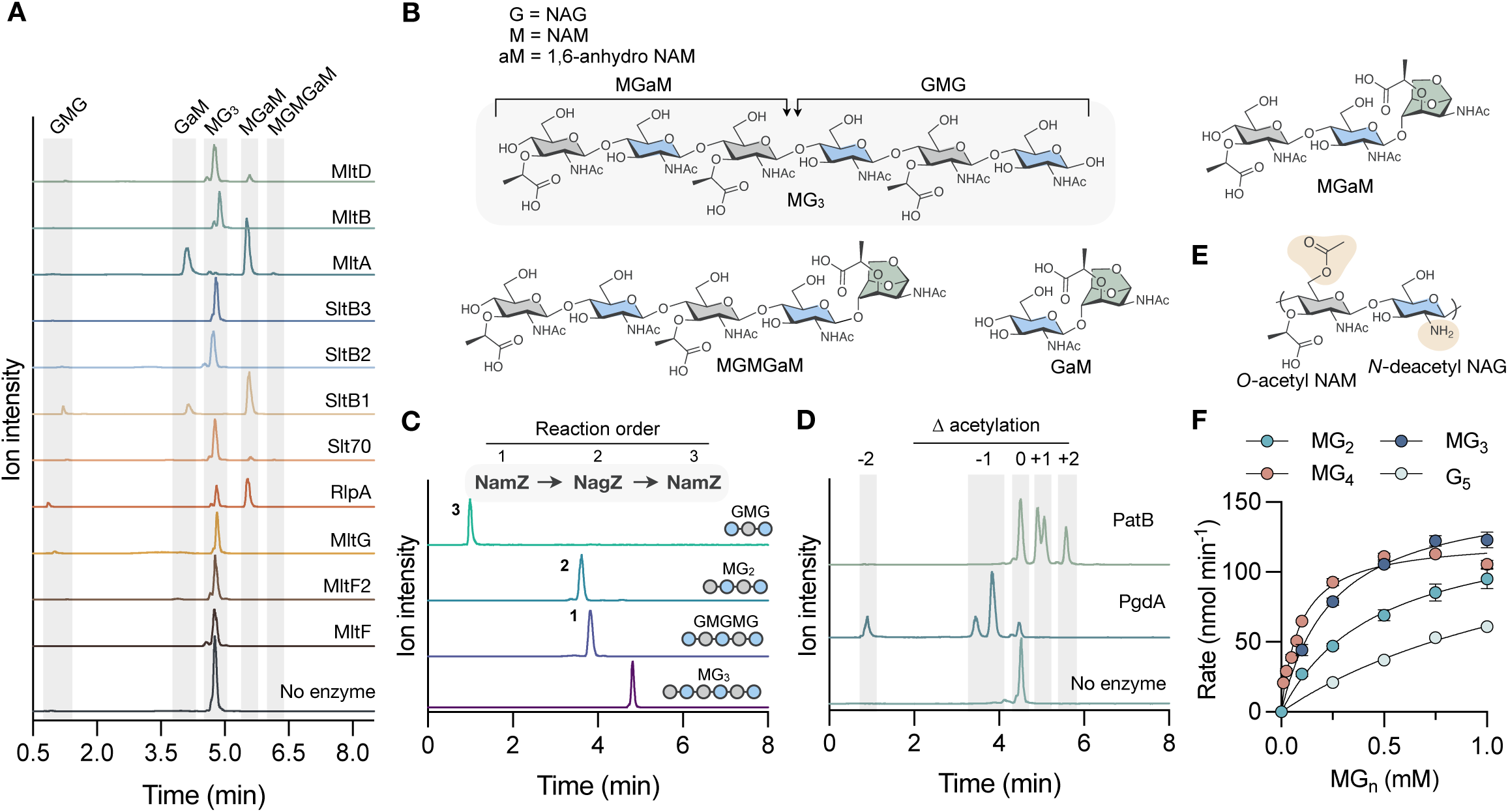
Enzymatic modification of defined denuded PG chains. *A,* LC-MS analysis of reaction mixtures with ten different LTs from *P. aeruginosa* and MG_3_. The proposed structures of the reaction products were determined by MS and represented as shaded bars by elution time. *B,* Schematic diagram of expected LT cleavage sites and structures of observed reaction products. *C,* LC-MS analysis of alternating exo-glycosidase activity using MG_3_ as the initial substrate (purple). Sequential cleavage by NamZ and NagZ is denoted by the reaction order. Representative glycan structures are depicted as interconnected circles, with blue and grey representing NAG and NAM, respectively. *D,* LC-MS analysis of changes in the extent of MG_3_ acetylation shown as shaded bars. *N-*deacetylation of NAG and O-acetylation of NAM were catalyzed by PgdA and PatB, respectively. *E,* Representative structure of O-acetylated and N-deacetylated glycan chains. *F*, Steady-state kinetic analysis of PatB-catalyzed O-acetylation of denuded PG glycans with varying degrees of polymerization using pNP-Ac as acetyl donor. Data represent the mean ± SD (n = 3).

As MG_3_ starts with a NAM residue at the non-reducing end, we next assessed whether exo-acting enzymes could sequentially hydrolyze the glycan chain. In line with this structural assignment, the exo-glucosaminidase NagZ did not cleave MG_3_, whereas the muramidase NamZ removed a single NAM residue to make the pentasaccharide GMGMG **(Fig. 3C and Fig. S5)**. Together, NamZ and NagZ acted sequentially to fully convert MG_3_ into NAM and NAG monosaccharides, consistent with previous studies using disaccharide fragments (**Fig. S5)**^26,27^. Because each residue is removed separately, using NamZ and NagZ in series cleaves one specific residue at a time **(Fig. 3C).** Together, these data demonstrate that denuded glycans are substrates for exo-acting hydrolases and a subset of LTs and that these enzymes can be applied as useful enzymatic tools for further tailoring defined glycan chains to create additional structural diversity.

### O-acetyltransferases and N-deacetylases modify denuded PG at variable positions throughout the chain

Having established that denuded glycans serve as substrates for diverse PG backbone-cleaving enzymes, we next tested whether they could also be used to monitor non-degradative glycan modification. To this end, we tested enzymes that modify glycan chain acetylation, including the NAG N-deacetylase PgdA of *Clostridioides difficile* and the NAM O-acetyltransferase PatB of *Campylobacter jejuni,* which remove or add acetyl groups to the PG backbone, respectively **(Fig. 3E)**^28,29^. We first tested if NAG could be deacetylated within the polymer. LC-MS analysis of reaction mixtures containing MG_3_ and PgdA revealed that >75% of MG_3_ substrate was consumed, and two products containing a single deacetylated NAG (−42 Da) at different positions and one product with two deacetylated NAG residues (−84 Da) **(Fig. 3D)**. We next analyzed the MG_3_ reaction with PatB using LC-MS to monitor O-acetylation with *p*-nitrophenyl acetate (pNP-Ac) as the acetyl donor. New species were detected with mass shifts consistent with the modification of one or two NAM residues (+42 and +84 Da, respectively), indicating that PatB can accept MG_3_ as a substrate and modify it at one or two distinct NAM positions **(Fig. 3D)**. Comparing the positional specificities of PgdA and PatB toward MG_3_ by MS/MS showed that both enzymes were restricted to modifying internal residues and did not modify the respective reducing or non-reducing ends **(Fig. S6 and S7)**. Together, these findings show that MG_3_ is a tractable substrate for revealing the position-dependent preferences of enzymes that modify the glycan chain. Moreover, the significant production of acetylated/deacetylated products by these enzymes represents yet another structural permutation of the defined denuded glycans that can be made, enabling interrogation of how glycan modifications influence enzyme recognition, substrate specificity, and PG interactions.

### Denuded PG is suitable for determining the chain length specificity of PG-modifying enzymes

Since we were able to purify MG oligomers of different lengths from a single batch of PG sacculi, we assessed their suitability for evaluating the length specificity of PG-active enzymes. To do this, we chose to assess the Michaelis-Menten kinetics of PG O-acetylation by PatB because (1) the transfer of O-acetyl groups from the pNP-Ac donor can be continuously monitored spectrophotometrically, and (2) the assay allows direct comparison of PatB kinetics with chitooligosaccharides, which serve as PG analogs for many PG-active enzymes^30,31^. Kinetic analyses of PatB revealed that the turnover rate for each DP was similar, centring around 21 s^-1^ **(Table 1).** However, PatB displayed a clear acceptor-length dependence, with a strong preference for MG_4_, as reflected by a *K*_m_ of 92 μM that increased in ∼2-fold increments as the substrate chain length decreased for each DP value **(Fig. 3F and Table 1)**. Compared with previous kinetic analyses of PatB using chitopentaose (G_5_; *K*_m_ = 1.6 mM), the *K*_m_ for O-acetyltransfer to MG_3_ (*K*_m_ = 0.25 mM), which is similar in length, was 6-fold lower **(Fig. 3F and Table 1)**^31^. The lower observed *K*_m_ with authentic PG chains likely reflects the enzyme’s higher affinity for NAM in these substrates. Taken together, these data indicate that although chitooligosaccharides are convenient, commercially available PG analogs, their structural simplicity limits their recognition by PatB and likely other PG-active enzymes. Denuded glycans are still simplified PG substrates because they lack stem peptides.

**Table 1.** Acceptor length dependence on the steady-state kinetic parameters of *C. jejuni* PatB.

| Acceptor | $k_{cat}$<br>(s<sup>-1</sup>) | $K_m$<br>(μM) | $k_{cat}/K_m$<br>(M<sup>-1</sup>•s<sup>-1</sup>) |
| --- | --- | --- | --- |
| MG<sub>2</sub> | 23.1 ± 1.3 | 473 ± 60 | 4.9 × 10<sup>4</sup> |
| MG<sub>3</sub> | 26.3 ± 0.07 | 250 ± 19 | 1.05 × 10<sup>5</sup> |
| MG<sub>4</sub> | 20.7 ± 0.4 | 92 ± 7 | 2.25 × 10<sup>5</sup> |

Activity assayed using 2 mM pNP-acetate in 50 mM sodium phosphate, pH 7.0 in triplicate

Nevertheless, they preserve the native glycan structure and provide a more biologically relevant substrate than chitooligosaccharides for defining the substrate-length dependence and kinetic parameters of PG-active enzymes.

### Synthesis of pNP-MG_2_ for the development of a continuous muramidase assay

The simplicity of pNP-based assays inspired us to pursue the synthesis of a PG derivative for colorimetric detection of glycan cleavage. In previous reports, chromogenic chitooligosaccharides were generated by attaching pNP to the C1 position of chitooligosaccharides for use as lysozyme substrates^32^. These substrates were prepared enzymatically using hen egg white lysozyme (HEWL) to catalyze the transglycosylation of pNP-NAG to NAG_5_ via exchange of the terminal non-reducing NAG residue. To favour transglycosylation, the reaction mixture contained 60% DMSO to reduce water activity and slow hydrolysis^33^. Using a similar approach, we also used HEWL to catalyze the exchange of the terminal NAG residue of MG_2_ in the presence of excess pNP-NAG in 60% DMSO **(Fig. 4A)**. MG_2_ was chosen because it contains only one possible position for pNP-NAG exchange. In the absence of pNP-NAG, HPLC analysis showed that MG_2_ is stable to HEWL-catalyzed hydrolysis. When a 5-fold excess of pNP-NAG was included in the reaction mixture, we found that transglycosylation was the preferred reaction at pH 4.5 **(Fig. 4B and Fig. S8)**. The synthesis of pNP-MG_2_ was then scaled up, and the product was isolated by HILIC, yielding 98% purity as judged by HPLC **(Fig. S9)**. Purified pNP-MG_2_ was then employed in reactions with MltA. HPLC-based assays showed that pNP-MG_2_ was cleaved into MGaM and pNP-NAG, with no other products formed **(Fig. 4C)**. This result confirmed that pNP-MG_2_ can be recognized as a substrate by MltA and that the production of pNP-NAG by MltA may be detected spectrophotometrically through pNP release by coupling the reaction to an exo-β-*N*-acetylglucosaminidase. Previous kinetic analysis of *B. subtilis* NagZ-catalyzed pNP-NAG hydrolysis reported *K*_m_ = 172 μM, *k*_cat_ = 9.9 s^-1^, and *k*_cat_/*K*_m_ = 5.8 × 10^4^ M^-1^ s^-1^, indicating that this enzyme turns over the substrate with high efficiency^34,35^. Thus, we chose to use this NagZ homolog in our coupled assay, as it should not limit spectrophotometric detection of pNP-NAG products from cleavage of pNP-MG_2_. To perform the coupled assay, reaction mixtures contained NagZ and MltA at a 5:1 molar ratio. Assays containing varying substrate concentrations were continuously monitored by spectrophotometry at λ = 405 nm. The rates of pNP production were linear and increased in a substrate concentration-dependent manner, indicating that the assay was suitable for kinetic studies (**Fig. 4D**). Cleavage of pNP-MG_2_ by MltA followed Michaelis-Menten kinetics and gave the kinetic parameters *K*_m_ = 190 ± 29 μM, *k*_cat_ = 0.026 ± 0.0014 s^-1^, and *k*_cat_/*K*_m_ = 135 M^-1^ s^-1^. Together, these results establish pNP-MG_2_ as a suitable substrate for continuous, coupled monitoring of LT activity for enzymes that can recognize denuded PG.

**Figure 4.**
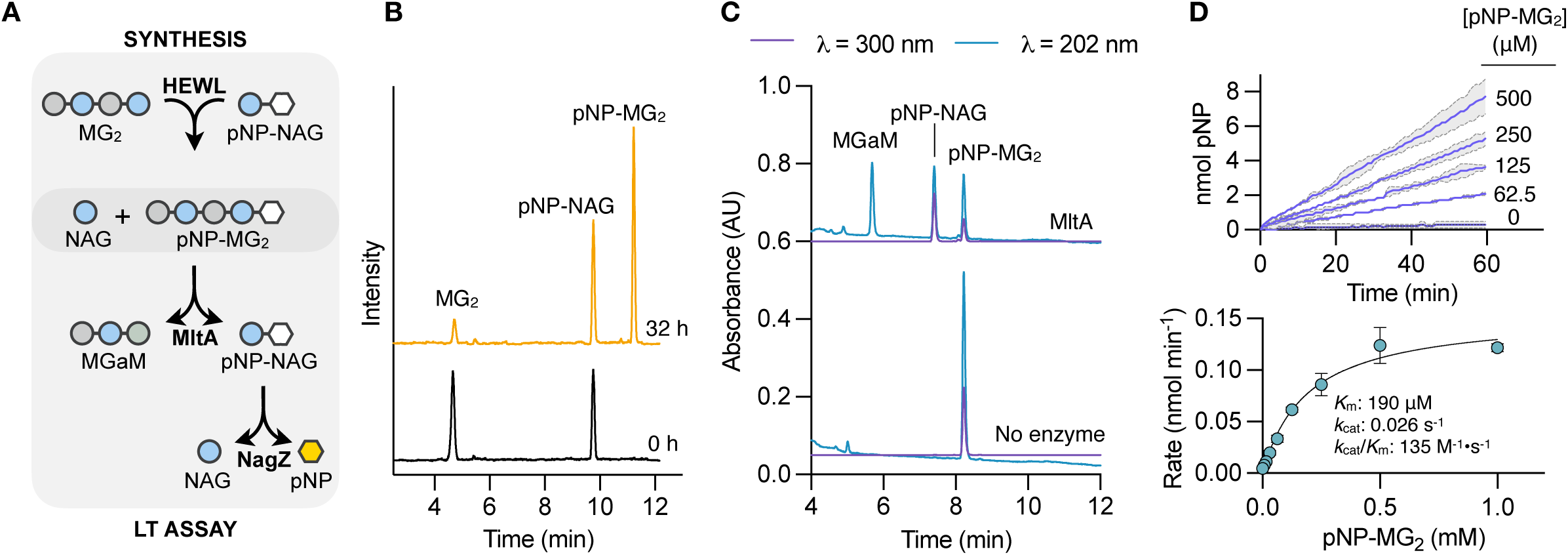
Development and validation of a colorimetric assay for lytic transglycosylases and muramidases. *A,* Reaction scheme for the production of pNP-MG_2_ and its subsequent cleavage as a NagZ-coupled colorimetric LT substrate. *B,* LC-MS analysis of HEWL-catalyzed transglycosylation using MG2 and pNP-NAG as acceptor and donor substrates, respectively. The transglycosylation reaction (32 h, yellow) was compared to the initial mixture of reactants (0 h, black). *C,* HPLC analysis of pNP-MG_2_ cleavage by MltA (top) compared to the untreated starting substrate (bottom). *D, Top,* Spectrophotometric time course of pNP-MG_2_ cleavage by MltA (1 μM) coupled to pNP-NAG hydrolysis by NagZ (5 μM). pNP release was monitored spectrophotometrically at 420 nm. Purple lines represent the mean ± SD (n = 2). SD is shown by the grey shaded area. *Bottom,* Michaelis-Menten kinetics of pNP-MG_2_ cleavage by MltA (1 μM) in 25 mM sodium acetate buffer, pH 6.5. Data represent the mean ± SD (n = 2).

## Discussion

In this study, we present an approach for producing, purifying, and diversifying defined PG glycan substrates from bacterial cell walls. By optimizing controlled digestion of PG sacculi isolated from staphylococci, this method provides direct access to denuded glycan chains of defined length. Moreover, it extends previous enzymatic digestion and fragment-isolation approaches by generating substantially longer oligosaccharides. Depending on the digestion conditions and amount of PG starting material used, purification by reversed-phase flash chromatography can yield hundreds of milligrams of highly pure denuded glycan chains. These glycan structures are ultimately suitable for *in vitro* studies of PG-active enzymes to facilitate the broader study of PG metabolism and support the discovery of new antimicrobial strategies.

Controlled digestion of staphylococcal PG reproducibly produced glycan chains from DP_2_ to DP_5_, with yields ranging from 0.5–4% of the cell wall dry weight depending on chain length and digestion conditions. This represents an important practical advance over prior enzymatic approaches that exhaustively digested *M. luteus* PG with lysozyme, which largely limited the major products to short glycan fragments of DP_1_ and DP_2_^18,20,21^. Moreover, our workflow outlines rapid chromatographic techniques for both the analysis and purification of longer glycan chains, offering a significant technical advantage over previous low-resolution, time-consuming size-exclusion chromatography methods^20^.

Another important feature of this workflow was the use of staphylococci as a source of PG. Compared with traditional *M. luteus* preparations, *S. carnosus* and *S. aureus* provided reliable starting material for large-scale cell-wall isolation, with *S. carnosus* offering the additional advantage of being a non-pathogenic, food-grade organism. Following removal of stem peptides, controlled digestion with LytD generated NAG-terminating glycan products with limited sequence heterogeneity, and individual oligosaccharides could be isolated to sufficient purity for biochemical assays. Production of longer oligosaccharides remains a limitation, as products larger than MG_5_ were obtained only in low yields because LytD rapidly converts longer chains into shorter products. Future optimization will therefore benefit from screening *N*-acetylglucosaminidases with distinct cleavage preferences or further tuning digestion conditions to preserve longer intermediates.

The terminal structure of PG chains reflects the enzymatic processes that synthesize, remodel, and degrade the cell wall. Many existing enzymatic and synthetic approaches generate glycans that terminate in reducing NAM residues, reflecting biosynthetic polarity or muramidase digestion. However, the terminal units of mature PG are notably more structurally variable across bacterial taxa. In diderm bacteria, such as *Escherichia coli*, PG chains terminate in 1,6-anhydro-NAM owing to the activity of lytic transglycosylases^4^. In contrast, in monoderm bacteria, such as *B. subtilis*, the terminal structures are expected to depend on the combined activities of muramidases, glucosaminidases, and lytic transglycosylases encoded in the genome^36,37^. The method outlined here provides access to a complementary class of defined PG glycans that terminate in reducing NAG residues. This topology is distinct from the NAM-terminated products generated by many existing enzymatic or synthetic approaches and expands the repertoire of defined PG substrates available for biochemical studies.

Denuded glycan chains specifically reflect products of amidase-processed PG. During cell division, amidases remove stem peptides from septal PG to promote daughter-cell separation and to define zones of new PG biosynthesis^38^. Therefore, the peptide-free glycan backbone is representative of remodeled septal PG of actively dividing cells. Given the biological context in which denuded PG is produced, the enzyme assays performed here illustrate both the utility and the limits of NAG-terminating denuded PG as a general substrate for PG-active enzymes. MG_3_ was accepted by several LTs and exo-acting glycosidases, demonstrating that the NAG-terminating glycan backbone alone is sufficient for productive binding for many enzymes. In cases where strong LT activity was observed (MltA, SltB1, and RlpA), these data were consistent with their known denuded PG activity and associations with the septal region of dividing cells^39–42^. The other LTs showed marginal (Slt70 and MltD) or no activity (MltF, MltF2, MltG, SltB2, SltB3, and MltB), indicating that stem peptides or other features of native PG remain important determinants of recognition for certain PG-active enzymes.

These observations complement previous work that first defined the activities of the eleven known LTs from *P. aeruginosa* using synthetic PG tetrasaccharides with the sequence NAG-NAM(pentapeptide)-NAG-NAM(pentapeptide)^24^. In that study, the presence of stem peptide and a terminal anhydro-NAM was important for recognition by all LTs, whereas tetrasaccharides terminating in NAM and/or lacking stem peptides were recognized by fewer enzymes. Notably, the denuded tetrasaccharide terminating in a reducing NAM was cleaved only by SltB1. These findings led to the conclusion that anhydro-NAM, the product of the LT reaction, is an important specificity determinant for some LTs, including MltA. This contrasts with our observations for the longer NAG-terminating substrate, MG_3_, which supported activity by MltA, RlpA, and SltB1. These differences indicate that the longer NAG-terminating denuded PG occupy a distinct glycan topology and broaden the range of available substrates for assessing the enzyme activity of PG-cleaving enzymes.

Thus, the denuded glycans described here are useful biochemical tools that aid in discriminating enzyme specificity. However, when the structural requirements for activity are unclear, they should be interpreted as simplified PG substructures rather than complete replacements for native PG. In this role, they serve as a bridge between heterogeneous cell-wall material, non-native chitooligosaccharides, and fully synthetic PG fragments. The PatB O-acetyltransferase assays further highlight this utility. Unlike chitooligosaccharides, the denuded glycan backbone enabled more accurate estimation of substrate length and positional specificity while remaining experimentally tractable.

Beyond their direct use as substrates, denuded glycans provide scaffolds for further diversification. Enzymes such as NamZ, NagZ, PgdA, and PatB can be used to tailor glycan length, terminal structure, and acetylation state, thereby creating additional, well-defined PG substructures from a common set of isolated glycans. We further explored this potential by generating a pNP derivative of PG via lysozyme-catalyzed transglycosylation. When coupled to NagZ, cleavage of this substrate enabled continuous spectrophotometric monitoring of lytic transglycosylase activity, establishing a new assay format based on a defined PG-derived substrate rather than a non-native chitooligosaccharide analog. The same logic can be extended to other PG-derived structures. Denuded glycans provide a useful scaffold for rebuilding more complex PG fragments, including structures in which defined stem peptides are reattached. This offers a practical advantage over total synthesis, in which constructing β(1,4)-glycosidic bonds in the PG backbone is often among the most challenging and inefficient steps ^43^. The significance of this bottleneck is illustrated by prior top-down efforts to prepare NAG-NAM structures from chitooligosaccharides, which use these natural polymeric templates to avoid multiple rounds of chemical glycosylation. Thus, the denuded glycans produced here are valuable not only as substrates but also as starting materials for the semi-synthesis of PG fragments, modified glycans, and chemical probes.

Overall, this work establishes a practical platform for accessing defined PG glycan substrates from bacterial cell walls. The method does not replace total synthesis, nor does it fully capture the structural complexity of native PG. Instead, it fills an important gap between heterogeneous cell-wall material, commercially available non-native analogs, and technically demanding synthetic substrates. By enabling the production, purification, enzymatic modification, and assay development of defined PG glycans, this approach should facilitate broader biochemical investigation of bacterial PG-active enzymes and proteins that recognize, process, or respond to PG-derived ligands.

## Acknowledgments

We thank members of the Center for Microbial Chemical Biology and the University of Waterloo Mass Spectrometry Facility for technical support and the use of mass spectrometry resources. We also extend our deepest gratitude to Gerry Wright for his early support of this project.

## Author contributions

B.G.E, G.D., and D.S. writing – original draft; B.G.E., G.D., A.C.A., C.V., D.S. validation; B.G.E., G.D., A.C.A., C.V., D.S. methodology; B.G.E., G.D., A.C.A., C.V., D.S. investigation; B.G.E, D.S. conceptualization; B.G.E., A.J.C., D.S. writing – review and editing; D.S. supervision; A.J.C. and D.S. resources; D.S. project administration; D.S. funding acquisition.

## Funding and additional information

This project was funded by the Natural Sciences and Engineering Research Council under the Discovery Grants RGPIN-2025-04548 to DS and RGPIN-2022-03986 to AJC, and Discovery Launch Supplement Award DGECR-2025-00426, and start-up funds from the University of Waterloo Department of Chemistry to D.S.

## Methods

### DNA manipulation and protein purification

Plasmids, constructs, and source DNA are summarized in **Table S1** of the supplementary methods. Briefly, gene fragments were PCR-amplified from chromosomal DNA using the appropriate primers and cloned into their respective plasmids via restriction sites included in the forward and reverse primers, respectively. The corresponding plasmid was assembled using T4 DNA ligase (NEB) according to the manufacturer’s protocol to yield the cloned gene. All assembled plasmids were verified by sequencing and restriction digestion.

The conditions for protein production are summarized in **Table S2** of the supplementary methods. Generally, plasmids were transformed into chemically competent *E. coli* expression strain using the heat-shock method. Overnight cultures of *E. coli* carrying the appropriate plasmid were diluted 1:100 into sterile liquid medium.

Production of PatB, MltA, MltB, SltB1, and Slt70 used super broth, and the production of MltD, MltF, MltF2, MltG, RlpA, SltB2, SltB3, and PgdA used LB. The batch cultures were grown at 37 °C with aeration to an optical density (at 600 nm) of 0.5-0.8, and induction was achieved by adding isopropyl β-D-1-thiogalactopyranoside (IPTG). Induction was continued depending on the protein being produced. Following induction, cell pellets were harvested by centrifugation (5,000 × g, 15 min, 4 °C) and stored at −20 °C.

For purification, cell pellets were resuspended in lysis buffer at a volume of 50 mL per L culture. DNase (1 mg), RNase (0.5 mg), and one cOmplete protease inhibitor cocktail tablet (Roche) were added to the cell suspensions before being disrupted by sonication at 4 °C. Unbroken cells were removed from the crude lysate by centrifugation (12,000 × g, 20 min, 4 °C). Insoluble debris was removed from the cleared lysate by centrifugation (28,000 × g, 30 min, 4 °C). The resulting soluble protein preparation was loaded onto 2 mL of cOmplete Ni purification resin (Roche, Mississauga, ON). Unless otherwise stated, the resin was washed three times in 10 mL of lysis buffer, then three times in 10 mL of wash buffer (lysis buffer plus 50 mM imidazole). The protein was eluted into elution buffer (lysis buffer with 300 mM imidazole) and was dialyzed against 50 mM HEPES, pH 7.5, 150 mM or 50 mM Tris, pH 7.5, 150 mM NaCl at 4 °C.

### Isolation of PG Sacculi and Liberation of Denuded Glycans

*S. carnosus* was grown for 24 hours in 21 L of LB broth (Lennox) at 37 °C with shaking at 250 rpm. Cells were then pelleted through centrifugation (5,000 × g for 20 minutes). Pellets were resuspended in Milli-Q water, mixed 1:1 with 8% (w/v) SDS solution to reach a final concentration of 4% (w/v) SDS, and boiled for 30 minutes while stirring to extract an insoluble PG product. SDS was washed out of the PG solutions by repeated centrifugation (12,000 × g for 10 minutes) and resuspension in warm Milli-Q water. Following the last centrifugation, pellets were resuspended in 25 mL of 0.1 M NaOH and left at 37°C for 4 hours to hydrolyze bulky groups attached to PG through phosphate links. Solutions were neutralized to pH 7.0 using 1:1 HCl:Milli-Q water. PG mesh was broken up through bead beating (15 rounds of 1 minute bead beating and 1 minute resting). Samples were vacuum filtered, and filtrates were collected and pooled together. Residual salts were removed from pooled filtrate through 2 rounds of centrifugation (12,000 × g for 6 minutes) and resuspension in 30 mL of phosphate-buffered saline buffer. To degrade any protein remaining in the sample, 1 mg of Pronase from *Streptomyces griseus* was added to the sample and incubated at 37 °C overnight. To inactivate the Pronase, the sample was boiled in an 80 °C water bath for 1 hour. Pronase, excess protein debris, and buffer salts were washed out of the sample through three rounds of centrifugation (12,000 × g for 10 minutes) and resuspension in Milli-Q water. The sample was lyophilized for 3 days to obtain dry yield. Dried PG was reconstituted in 25 mM ammonium acetate buffer (pH 7.5) and 20 μg/mL recombinant AmiA from *S.aureus* was added to cleave stem peptides. The sample was incubated at 37 °C for 24 hours until turbidity was constant, indicating completion of stem peptide cleavage. Acetic acid was added dropwise to the sample until it reached pH 3.0 to acidify cleaved stem peptides (holding a positive charge) and denuded glycans (holding a negative charge). Soluble stem peptides and denuded glycans were separated from excess insoluble debris through centrifugation (12,000 × g for 90 minutes). The supernatant was passed through a 5 mL cation exchange column (AmberChrom 50WX4 Hydrogen resin), and peptide separation was monitored using acidic ninhydrin while spotting on a TLC plate. Denuded glycan filtrate was lyophilized for 3 days to obtain pure denuded glycan yield.

### Limited Digestion and Purification of Denuded Glycans

Denuded glycans were reconstituted in a minimal amount of 25 mM ammonium acetate buffer (pH 7.5). The pH of the solution was verified and titrated as necessary with 10% ammonium hydroxide. Various concentrations of LytD were tested in a small-scale reaction to determine optimal digestion parameters. For the large-scale reaction, 60 μg/mL of LytD was added and the reaction was incubated at room temperature for 30 minutes. Glycan cleavage was monitored through HPLC (Agilent 1100 series) using a Luna 5 μm HILIC column at 205 nm nm using the following method: Solvent A: water + 0.1% formic acid, solvent B: acetonitrile + 0.1 % formic acid; flow rate: 1 mL/min, gradient: 0 - 1 min 80% B, 1 – 15 min to 50% B, 15 – 15.1 min to 80% B, 15.1 – 20 min at 80%. Following HPLC chromatogram verification of MG_2_ as the dominant fragment after 2 hours, the digestion mixture was quenched with 0.1% FA and boiled at 80 °C for 10 minutes.

The polymer mixture post-digestion was centrifuged (21,100 × g for 3 minutes) and the supernatant was syringe-filtered through a 0.45 μm filter. Glycan oligosaccharides were separated using flash chromatography (RediSep Rf reverse phase C18aq column) through a stepwise gradient of 3 C.V. of 0% to 5% ACN + 0.1% FA. The column was activated with 10 C.V. of 50% ACN + 0.1% FA and equilibrated with 10 C.V. of Milli-Q water + 0.1% FA. Fractions were checked for purity and predominant polymer using mass spectrometry (Thermo Scientific LTQ-XL Linear Ion Trap) through electrospray ionization. Resulting fractions containing pure MG_2_ were pooled together, concentrated to 5 mL, then lyophilized to obtain a final yield of pure MG_2_.

### Hydrophilic interaction chromatography and MS/MS analysis of glycans

Purity assessment of the final purified glycans was assessed by HPLC (Agilent 1100 series) using a Phenomenex Luna 5 μm HILIC 200 Å column (4.6 × 100 mm) at 205 nm using the following method: Solvent A: water + 0.1% formic acid, solvent B: acetonitrile + 0.1 % formic acid; flow rate: 1 mL/min, gradient: 0 - 1 min 80% B, 1 - 15 min to 50% B, 15 – 15.1 min to 80% B, 15.1 – 20 min at 80% B; wavelength: 205 nm. The identity of the samples was verified using mass spectrometry (Thermo Scientific LTQ-XL Linear Ion Trap) with electrospray ionization. Prior to mass spectrometry analysis, samples were reduced with the following method: samples were mixed 1:1 with 500 mM sodium borate buffer containing sodium borohydride (1 % (w/v)) and left to reduce (20 min, RT), and excess borohydride was destroyed with 1% formic acid at the end of incubation. The samples were desalted by HPLC using a C8 column, directly infused into an LTQ-XL Linear Ion Trap (Thermo Scientific), and analyzed in positive ion mode.

### LC-MS-based assays

LT-mediated glycan cleavage reactions with LTs were carried out under buffer conditions described previously^24^. Briefly, each enzyme (1 μM) was incubated with MG_3_ (200 μM) in 20 mM HEPES, 0.1 M NaCl, pH 7 at 37 °C for 1 hour. Reactions with NagZ (1 μM), NamZ (1 μM), or both were performed with MG_3_ (200 μM) in PBS, pH 7.5 at 37 °C for 1 hour. O-acetylation reactions with PatB (1 μM) were performed with MG_3_ (200 μM) and 1 mM pNP-acetate in 25mM HEPES, pH 7.5 at 37 °C for 16 hours.

Deacetylation reactions with PgdA (1 μM) were performed in 25mM HEPES, pH 7.5at 37 °C for 16 hours using MG_3_ (200 μM). Following incubation, all enzymes were heat-inactivated (100 °C, 5 mins), and the samples were mixed 1:1 with 500 mM sodium borate buffer containing sodium borohydride (1 % (w/v)) and left to reduce (20 min, RT), and excess borohydride was destroyed with 1% formic acid at the end of incubation. Samples were analyzed by LC-MS using an Agilent Zorbax 1.9 μm XDB-C8 column (2.1 mm × 100 mm) column on a Vanquish Core UPLC in line with an LTQ-Linear Ion Trap (Thermo Scientific) using the following method: Solvent A: water + 0.1% formic acid, solvent B: acetonitrile + 0.1% formic acid; flow rate: 0.4 mL/min, gradient: 0 min 1% B, 0 - 12 min to 25% B; detection: MS in negative ion mode.

### Colorimetric O-acetyltransferase assay and enzyme kinetics

The assay for PG *O*-acetyltransferase was used to measure the activity of recombinant PatB as described previously^30,31^. PatB (1 µM) was mixed with 2 mM *p*-nitrophenyl acetate (prepared in 100% DMSO) and denuded PG glycans at the specified concentration in 50 mM sodium phosphate pH 7.0. The final DMSO content of assays was 5% (v/v). Reactions (100 µL) were performed in triplicate at 37 °C and the absorbance at 405 nm was monitored in an imaging microplate reader (BioTek, Winooski, VT) every 60 s for 15 min. Model fitting and kinetic parameters were obtained with GraphPad Prism (v10.2.1).

### Transglycosylation of pNP-NAG to MG_2_

Reactions were prepared with the following final concentrations: 10 mM MG_2_ in Milli-Q water, 60 mM pNP-GlcNAc in DMSO, and 174 µM HEWL in 50 mM sodium acetate buffer (pH 4.0, 4.5, or 5.0). A separate reaction was prepared at pH 5.0 with no HEWL as a negative control. Reactions were incubated at 30 °C and aliquots taken daily. Prior to monitoring the reactions, the sample aliquots were mixed 1:1 with 0.5 M sodium borate buffer (pH 9.0) and boiled at 80 °C for 10 minutes. After boiling, aliquots were reduced with a minimal amount of sodium borohydride powder and incubated at RT for 20 minutes.

Reduction was quenched with 2 µL of formic acid. Aliquots were then diluted 1:5 with Milli-Q water, centrifuged (13,800 × g), and the supernatant was monitored by HPLC (Agilent 1100 Series) using an Agilent Zorbax 1.9 μm XDB-C8 column (2.1 mm × 100 mm), with detection at wavelengths of 205 nm and 240 nm using the following method: Solvent A: water + 0.1% formic acid, solvent B: acetonitrile + 0.1 % formic acid; flow rate: 0.5 mL/min, gradient: 0 - 1 min 1.5% B, 1 - 13 min to 40% B, 13 – 13.2 min to 50% B, 13.2 – 13.7 min at 50% B, 13.7 – 14 min to 1.5% B, 14 – 16 min at 1.5% B. Chromatograms were compared to determine the optimal pH for the large-scale transglycosylation reaction using pNP-GlcNAc. Large-scale reactions were carried out in mixtures (100 µL) with 5 mM MG_2_ (5 mg), 60 mM pNP-NAG (20.5 mg), 60% (v/v) DMSO, in 16 mM sodium acetate buffer, pH 4.0, at 30 °C for 48 h.

Following incubation, the starting material and products were precipitated with acetonitrile at a final concentration of 90%. The pellet was next dissolved in 250 μL water and mixed 1:1 with acetonitrile before purification. pNP-MG_2_ was isolated by HPLC with a semi-preparative Phenomenex Luna 5 μm HILIC 200 Å column (10 mm × 250 mm) column using the following method: Solvent A: water + 0.1% formic acid, solvent B: acetonitrile + 0.1 % formic acid; flow rate: 5 mL/min, gradient: 0 - 1 min 90% B, 1 - 20 min to 70% B, 20 – 20.1 min to 90% B, 20.1 – 26 min at 90% B; wavelength: 205 nm. This afforded pNP-MG_2_ in 14% yield and enabled recovery of 46% of the unused starting MG_2_.

### HPLC and coupled colorimetric detection of pNP-MG_2_ cleavage by MltA

To verify MltA cleavage of pNP-MG_2_, a reaction was conducted as follows: 100 µM pNP-MG_2_ reconstituted in Milli-Q water, 10 mM magnesium chloride reconstituted in Milli-Q water, 1 µM MltA in 25 mM HEPES, pH 6.5. A negative control was prepared in the same manner, excluding MltA. Reactions were incubated at 37 °C for 90 minutes. Aliquots of the reaction were analyzed by HPLC (Agilent 1100 Series) using an Agilent Zorbax 1.9 μm XDB-C8 column (2.1 mm × 100 mm), monitored at wavelengths of 202 nm and 300 nm. Upon confirmation of the production of pNP-NAG via HPLC, colorimetric detection of pNP-MG_2_ cleavage by sMltA was assessed using reaction mixtures (100 µL) containing varying concentrations of pNP-MG_2_ (ranging from 1 mM to 0 mM dissolved in Milli-Q water). MltA (1 µM) and NagZ (5 µM) were mixed with the substrate 50 mM HEPES, 10 mM magnesium chloride, pH 6.5. Production of pNP was monitored spectrophotometrically in duplicate at 405 nm using an imaging microplate reader (BioTek Powerwave XS) in 49 s intervals over one hour at ambient temperature (∼23°C). Model fitting and kinetic parameters were obtained with GraphPad Prism (v10.2.1).

## SUPPLEMENTAL FIGURE LEGENDS

**Figure S1. Optimization of denuded PG digestion conditions by varying LytD concentration.** Denuded glycan chains (30 mg/mL) obtained from AmiA digestion of *S. carnosus* cell walls were incubated with varying concentrations of LytD. Reaction mixtures were buffered with 25 mM ammonium acetate, pH 7.5, and were incubated for 30 min at room temperature. Following incubation, the reactions were quenched by mixing the samples 1:1 with acetonitrile containing 0.1% formic acid, and then analyzed by HILIC. Peak numbering represents the degree of polymerization per MG disaccharide unit. Letters (a and b) denote separated anomers.

**Figure S2. MS/MS analysis of defined denuded PG.** Reduced denuded glycan chains were dissolved in 50% ACN, 0.1% formic acid and analyzed in positive ion mode with an LTQ-XL Linear Ion Trap MS. Collision-induced dissociation was performed at an amplitude of 35. Fragmentation patterns correspond to the MS/MS spectrum shown above it.

**Figure S3. MS/MS analysis of WTA-linked denuded PG.** Unreduced denuded glycan chains were dissolved in 50% ACN, 0.1% formic acid and analyzed in negative ion mode with an LTQ-XL Linear Ion Trap MS. Collision-induced dissociation was performed at an amplitude of 35.

**Figure S4. HILIC analysis of denuded glycans purified by preparative reversed-phase flash chromatography.** Purified denuded glycan chains (2 mM) in 50% ACN, 0.1% formic acid were analyzed by HILIC with an Agilent 1100 HPLC system using a linear gradient from 80% ACN, 0.1% formic acid in water to 50% ACN, 0.1% formic acid in water over 15 min. Absorbance was monitored at a wavelength of 202 nm.

**Figure S5. LC-MS analysis of MG3 treated with NagZ and NamZ.** Reaction mixtures containing PBS (pH 7.4), purified MG3 (0.1 mM), and the enzyme were incubated for 1 hour at 37 °C. Each reaction contained NagZ or NamZ alone, or together at a final concentration of 1 μM. Reactions were analyzed by LC-MS fitted with an Agilent ZORBAX XDB-C8 column in negative ion mode.

**Figure S6. MS/MS analysis of denuded PG O-acetylated by PatB.** Reduced denuded glycan chains were dissolved in 50% ACN, 0.1% formic acid and analyzed in positive ion mode with an LTQ-XL Linear Ion Trap MS. Collision-induced dissociation was performed at an amplitude of 35. Fragmentation patterns correspond to the MS/MS spectrum shown above it.

**Figure S7. MS/MS analysis of denuded PG deacetylated by PgdA.** Reduced denuded glycan chains were dissolved in 50% ACN, 0.1% formic acid and analyzed in positive ion mode with an LTQ-XL Linear Ion Trap MS. Collision-induced dissociation was performed at an amplitude of 35. Fragmentation patterns correspond to the MS/MS spectrum shown above it.

**Figure S8. Optimization of HEWL-catalyzed transglycosylation of pNP-NAG to MG_2_.** Reaction mixtures containing 60% DMSO in water with 10 mM MG_2_, 60 mM pNP-NAG, 174 μM HEWL, and sodium acetate buffer (pH 4.0, 4.5, or 5) were incubated for 72 hours at 30 °C. Reactions were analyzed by HPLC using an Agilent ZORBAX XDB-C8 column, monitoring UV absorption at a wavelength of 205 nm.

**Figure S9. Purification and structural characterization of pNP-MG_2_.** *A*, Semi-preparative HILIC of pNP-MG_2_. The HPLC trace was recorded at a wavelength of 205 nm. *B,* Collision-induced dissociation MS/MS of pNP-MG_2_. Fragmentation was performed at an amplitude of 35, and the key fragment ions are labelled on the structure accordingly.

