## Supplementary Figures for "Denuded peptidoglycan oligosaccharides enable the biochemical investigation of bacterial cell wall recognition, modification, and degradation"

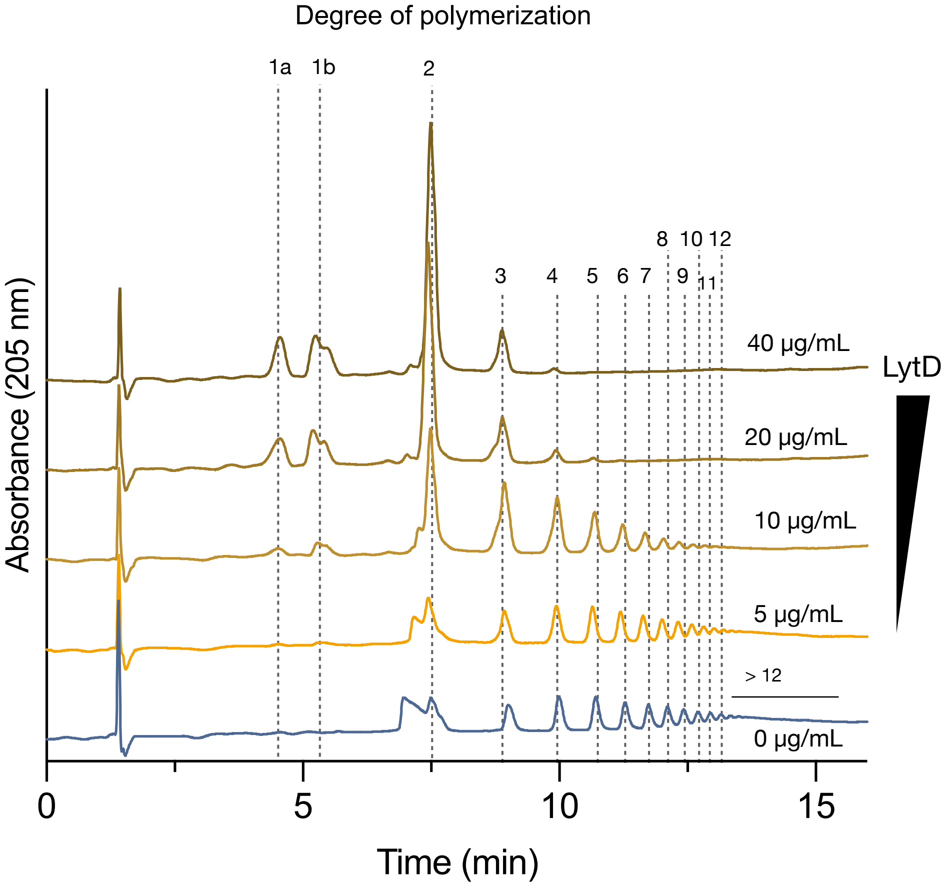


**Figure S1. Optimization of denuded PG digestion conditions by varying LytD concentration.** Denuded glycan chains (30 mg/mL) obtained from AmiA digestion of *S. carnosus* cell walls were incubated with varying concentrations of LytD. Reaction mixtures were buffered with 25 mM ammonium acetate, pH 7.5, and were incubated for 30 min at room temperature. Following incubation, the reactions were quenched by mixing the samples 1:1 with acetonitrile containing 0.1% formic acid, and then analyzed by HILIC. Peak numbering represents the degree of polymerization per MG disaccharide unit. Letters (a and b) denote separated anomers.

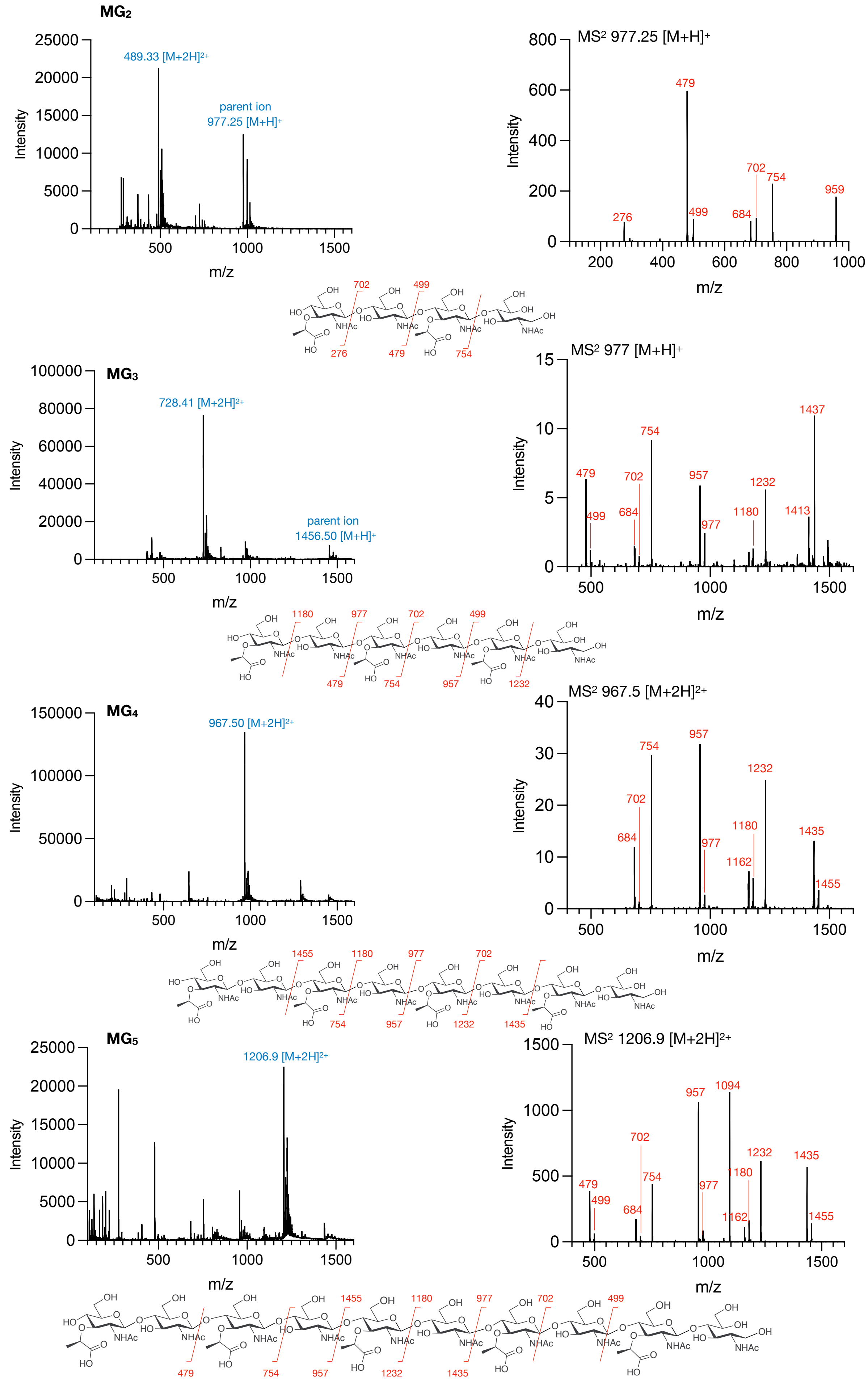


**Figure S2. MS/MS analysis of defined denuded PG.** Reduced denuded glycan chains were dissolved in 50% ACN, 0.1% formic acid and analyzed in positive ion mode with an LTQ-XL Linear Ion Trap MS. Collision-induced dissociation was performed at an amplitude of 35. Fragmentation patterns correspond to the MS/MS spectrum shown above it.

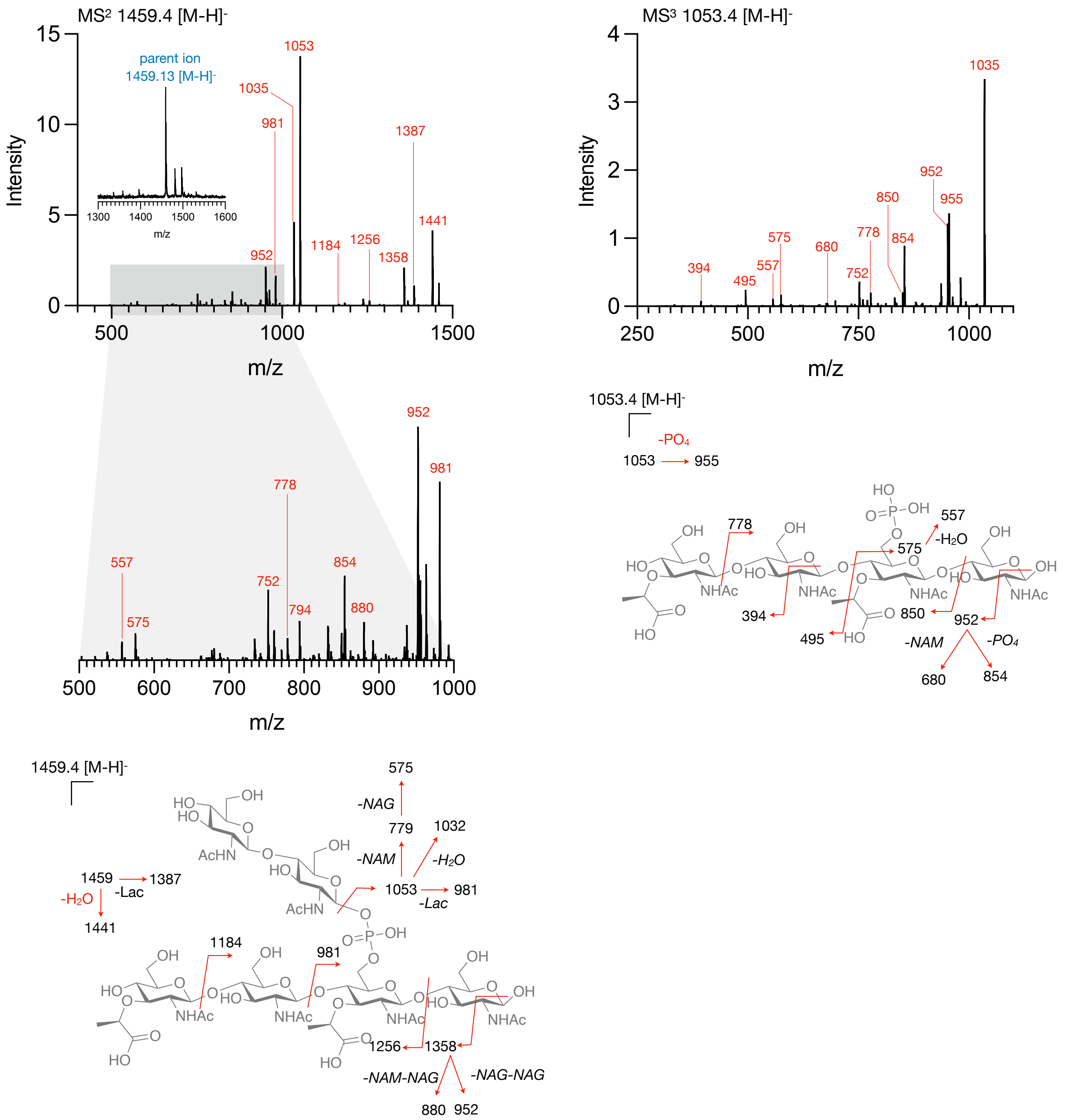


**Figure S3. MS/MS analysis of WTA-linked denuded PG.** Unreduced denuded glycan chains were dissolved in 50% ACN, 0.1% formic acid and analyzed in negative ion mode with an LTQ-XL Linear Ion Trap MS. Collision-induced dissociation was performed at an amplitude of 35.

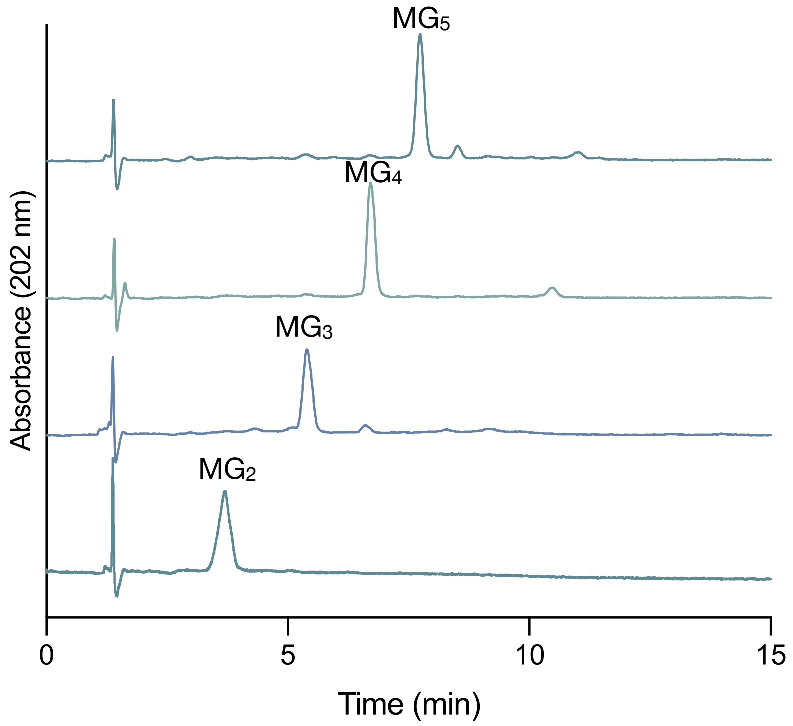


**Figure S4. HILC analysis of denuded glycans purified by preparative reversed-phase flash chromatography.** Purified denuded glycan chains (2 mM) in 50% ACN, 0.1% formic acid were analyzed by HILIC with an Agilent 1100 HPLC system using a linear gradient from 80% ACN, 0.1% formic acid in water to 50% ACN, 0.1 formic acid in water over 15 mins. Absorbance was monitored at a wavelength of 202 nm.

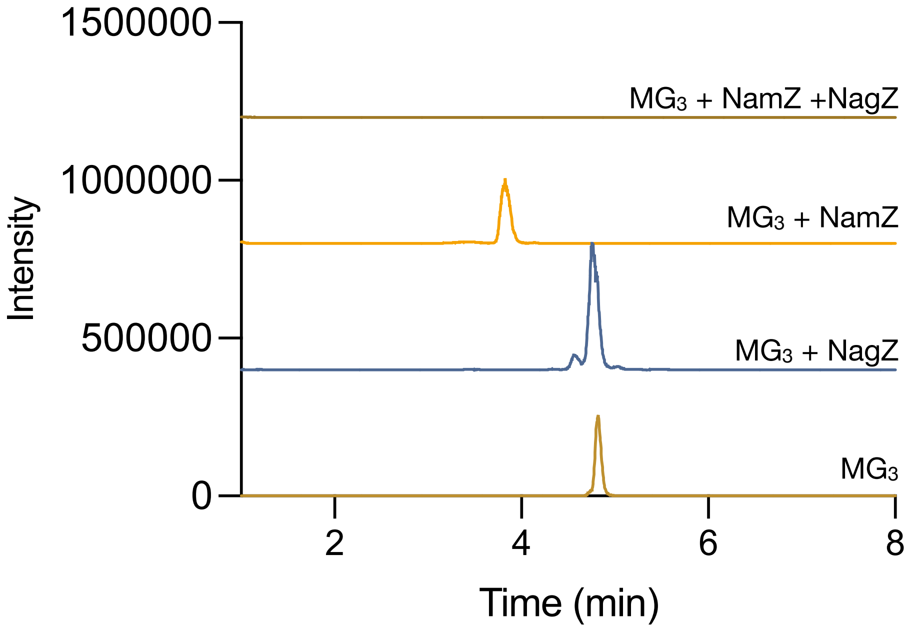


**Figure S5. LC-MS analysis of $MG_3$ treated with NagZ and NamZ.** Reaction mixtures containing PBS (pH 7.4), purified MG3 (0.1 mM), and the enzyme were incubated for 1 hour at 37 °C. Each reaction contained NagZ or NamZ alone, or together at a final concentration of 1 µM. Reactions were analyzed by LC-MS fitted with an Agilent ZORBAX XDB-C8 column in negative ion mode.

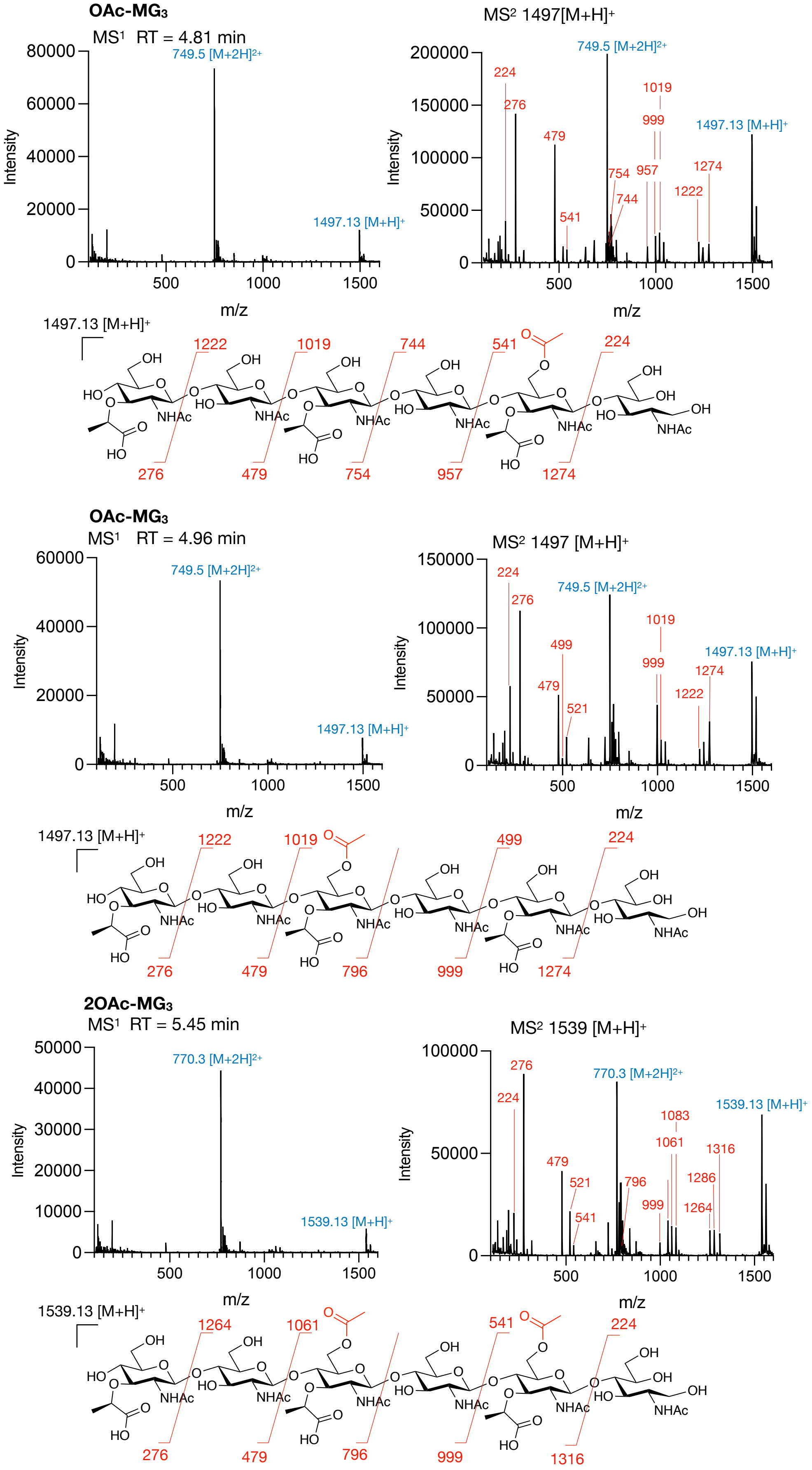


**Figure S6. MS/MS analysis of denuded PG O-acetylated by PatB.** Reduced denuded glycan chains were dissolved in 50% ACN, 0.1% formic acid and analyzed in positive ion mode with an LTQ-XL Linear Ion Trap MS. Collision-induced dissociation was performed at an amplitude of 35. Fragmentation patterns correspond to the MS/MS spectrum shown above it.

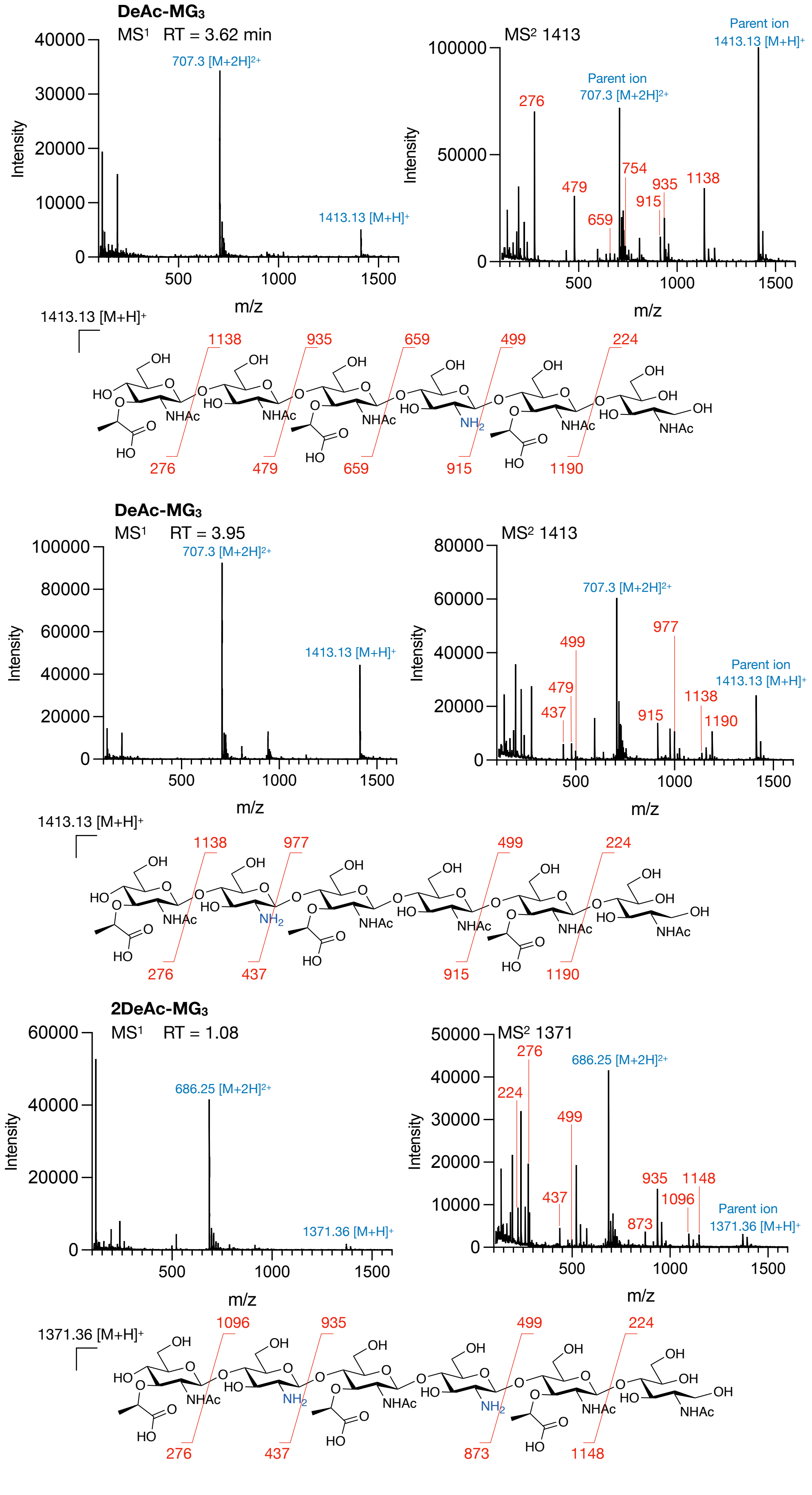


**Figure S7. MS/MS analysis of denuded PG de-acetylated by PgdA.** Reduced denuded glycan chains were dissolved in 50% ACN, 0.1% formic acid and analyzed in positive ion mode with an LTQ-XL Linear Ion Trap MS. Collision-induced dissociation was performed at an amplitude of 35. Fragmentation patterns correspond to the MS/MS spectrum shown above it.

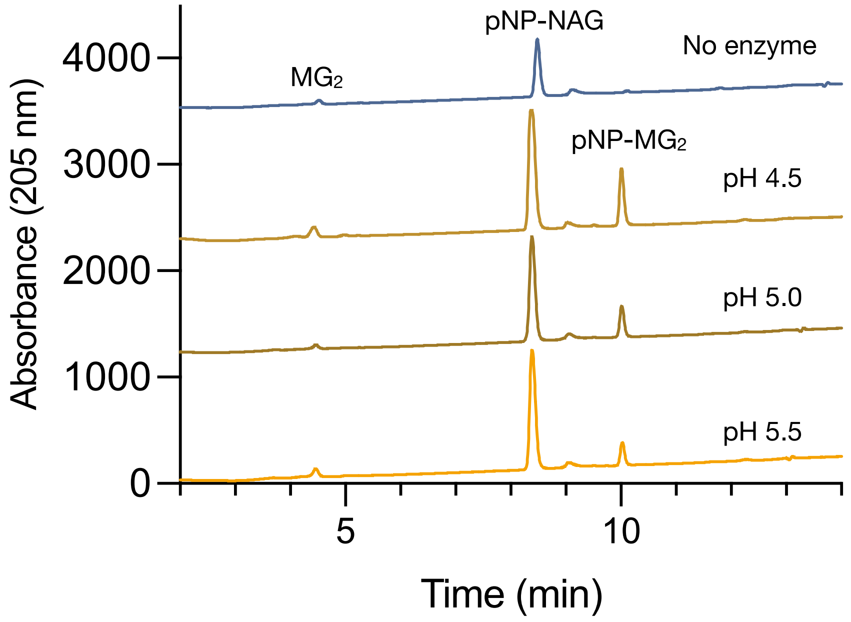


**Figure S8. Optimization of HEWL-catalyzed transglycosylation of pNP-NAG to $MG_2$.** Reaction mixtures containing 60% DMSO in water with 10 mM MG2, 60 mM pNP-NAG, 174 µM HEWL, and sodium acetate buffer (pH 4.0, 4.5, or 5) were incubated for 72 hours at 30ºC. Reactions were analyzed by HPLC using an Agilent ZORBAX XDB-C8 column, monitoring UV absorption at a wavelength of 205 nm.

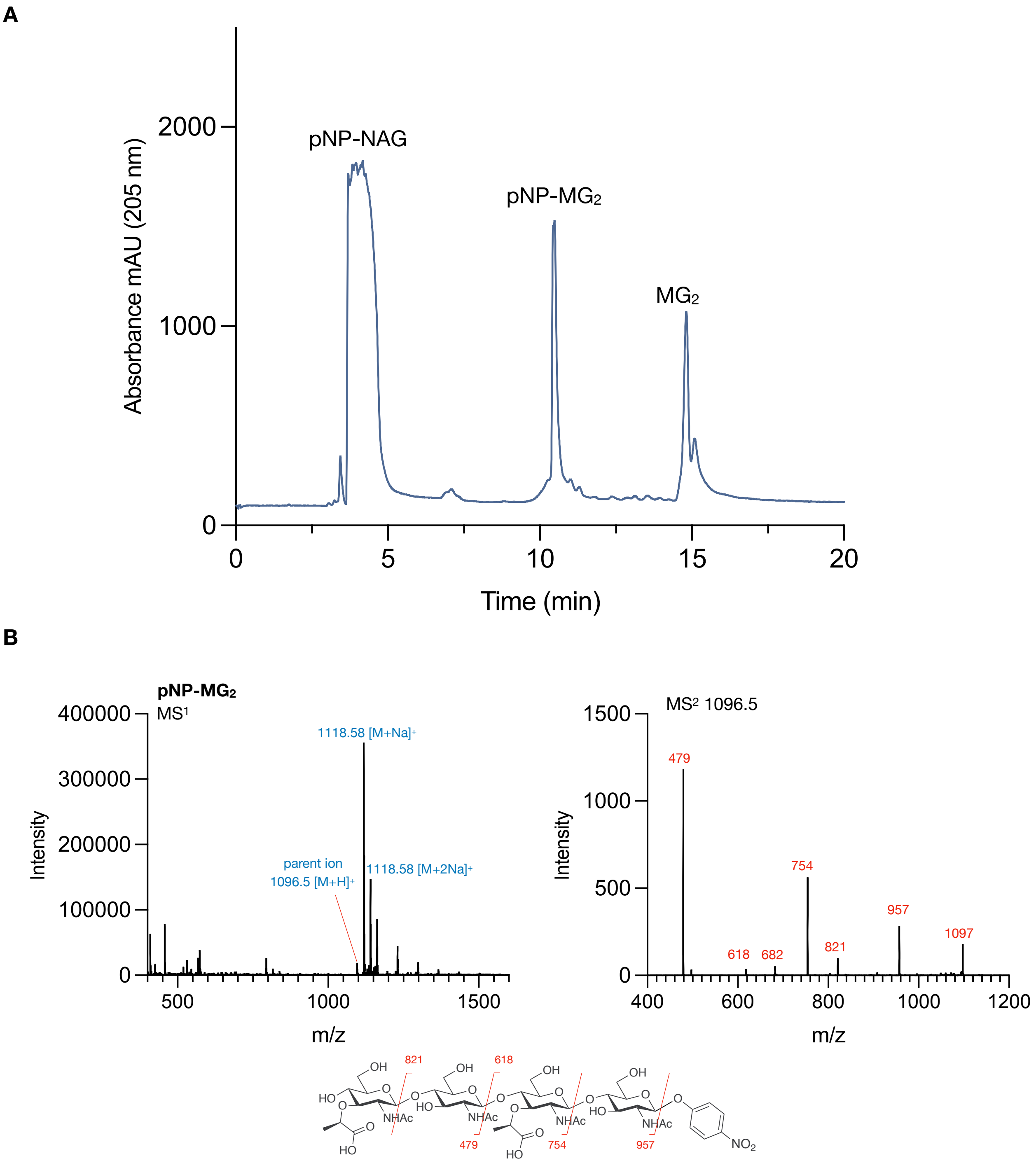


**Figure S9. Purification and structural characterization of pNP-MG$_2$.** *A*, Semi-preparative HILIC of pNP-MG$_2$. The HPLC trace was recorded at a wavelength of 205 nm. *B,* Collision-induced dissociation MS/MS of pNP-MG$_2$. Fragmentation was performed at an amplitude of 35, and the key fragment ions are labelled on the structure accordingly.
