## Supplementary methods for "Denuded peptidoglycan oligosaccharides enable the biochemical investigation of bacterial cell wall recognition, modification, and degradation"

| Table S1 |  |  |  |  |
| --- | --- | --- | --- | --- |
| Gene | Plasmid name | Plasmid backbone | Details | Expression strain |
| sMltA (PA1222) | pACCV-4 | pET28a(+) | truncated aa 1-25 | E. coli BL21(DE3) Star |
| sMltB (PA4444) | pNBAC54-1 | pET30a(+) | Ref: <https://pubs.acs.org/doi/10.1021/bi011833k> | E. coli BL21(DE3) pLysS |
| sMltD (PA1812) | pACCV-32 | pET28a(+) | truncated aa 1-34 | E. coli BL21(DE3) Star |
| sMltF (PA3764) | pACCV-21 | pET28a(+) | truncated aa 1-38 | E. coli SHuffle T7 |
| sMltF2 (PA2865) | pACCV-31 | pET28a(+) | truncated aa 1-21 | E. coli BL21(DE3) Star |
| sMltG (PA2963) | pACCV-37 | pBAD-SUMO | truncated aa 1-24 | E. coli BL21(DE3) Star |
| sRlpA (PA4000) | pACCV-33 | pET28a(+) | truncated aa 1-29 | E. coli BL21(DE3) Star |
| Slt70 (PA3020) | pACCV-47 | pET28a(+) | truncated aa 1-25 | E. coli BL21(DE3) Star |
| SltB1 (PA4001) | pNBAC258-2 | pET30a(+) | Ref: <https://pubs.acs.org/doi/10.1021/bi011833k> | E. coli BL21(DE3) pLysS |
| SltB2 (PA1171) | pACCV-29 | pET28a(+) | truncated aa 1-18 | E. coli BL21(DE3) Star |
| SltB3 (PA3992) | pACCV-28 | pET28a(+) | Truncated aa 1-32 | E. coli BL21(DE3) Star |

Growth medium - sMltA, sMltB, SltB1, Slt70 - super broth

- sMltD, sMltF, sMltF2, sMltG, sRlpA, SltB2, SltB3 - LB

Cells were grown aerobically at 37°C until an OD_600_ of 0.6-0.8 was reached.

Induction:

| Table S2 |  |  |  |
| --- | --- | --- | --- |
|  | Temp | Working [IPTG] | Duration |
| sMltA | 37°C | 0.1 mM | 16 h |
| sMltB |  | <https://pubs.acs.org/doi/10.1021/bi011833k> |  |
| sMltD | 15°C | 0.1 mM | 16 h |
| sMltF | 30°C | 1 uM | 16 h |
| sMltF2 | 15°C | 0.1 mM | 16 h |
| sRlpA | 15°C | 0.1 mM | 16 h |
| Slt70 | 15°C | 10 uM | 16 h |
| SltB1 |  | <https://pubs.acs.org/doi/10.1021/bi011833k> |  |
| SltB2 | 15°C | 0.4 mM | 16 h |
| SltB3 | 15°C | 1 uM | 16 h |

| sMltG | 15°C | 0.2% arabinose | 16 h |
| --- | --- | --- | --- |

Purification

Cell pellets from 1 L cultures were resuspended in lysis buffer containing 1 cOmplete™, Mini, EDTA-free Protease Inhibitor Cocktail tablet (Roche). Cells were lysed by sonication.

IMAC was performed in gravity columns packed with Roche cOmplete His-Tag Purification Resin.

sMltA, sMltD, sMltF2, sRlpA

Lysis buffer (L1): 50 mM Tris pH 8.0, 300 mM NaCl, 10% glycerol, 0.05% Brij-35, 5 mM EDTA, 20 mM imidazole

Elution buffer (E1): L1 + 50-1000 mM imidazole

sMltF, SltB2, SltB3

Lysis buffer: 50 mM sodium phosphate pH 8.0, 300 mM NaCl, 10 % glycerol

Wash buffer 1: L1 + 0.1% Triton X-100 pH 7.0

Elution buffer 1: L1 + 0.1% Triton X-100 pH 5.0

Elution buffer 2: L1 + 0.1% Triton X-100 pH 4.5

MltG purification

Lysis buffer (L1): 50 mM Tris pH 8.0, 300 mM NaCl, 10% glycerol, 0.05% Brij-35, 5 mM EDTA, 20 mM imidazole

Following lysis, cell-free lysate was applied to a gravity column packed with Roche cOmplete His-Tag Purification Resin. Washes were performed with 50 mL L1 followed by 50 mL L1 containing no NaCl. After washing, 10 mL L1 was added, then an aliquot of SUMO protease. This was incubated at room temperature for 1 h with gentle nutation. Flowthrough was collected, concentrated and loaded onto a HiLoad^TM^ 16/600 Superdex^TM^ 200 pg column (Cytiva). Size-exclusion chromatography was performed in 10 mM sodium acetate pH 5.0, 10 mM MgCl_2_, 10% glycerol.

Slt70 purification

Lysis buffer: 50 mM sodium phosphate pH 8.0, 300 mM NaCl

Wash buffer 1: lysis buffer pH 7.0

Wash buffer 2: lysis buffer pH 6.5

Wash buffer 3: lysis buffer pH 6.0

Elution buffer 1: lysis buffer pH 5.0

Elution buffer 2: lysis buffer pH 4.5

Elution buffer 3: 250 mM sodium phosphate, 300 mM NaCl pH 8.0

IMAC was performed in a gravity column packed with TALON® Metal Affinity Resin (Takara Bio). For the final step, E2 was eluted into E3 to prevent protein precipitation. Dialysis was subsequently performed in 10 mM sodium acetate pH 4.0 , 10 mM MgCl_2_, 10 % glycerol.
